# Two-color live-cell STED nanoscopy by click labeling with cell-permeable fluorophores

**DOI:** 10.1101/2022.09.11.507450

**Authors:** Carola Gregor, Florian Grimm, Jasmin Rehman, Christian A. Wurm, Alexander Egner

## Abstract

STED nanoscopy allows for the direct observation of dynamic processes in living cells and tissues with diffraction-unlimited resolution. Although fluorescent proteins can be used for STED imaging, these labels are often outperformed in photostability by organic fluorescent dyes. This feature is especially crucial for time-lapse imaging. Unlike fluorescent proteins, organic fluorophores cannot be genetically fused to a target protein but require different labeling strategies. To achieve simultaneous imaging of more than one protein in the interior of the cell with organic fluorophores, bioorthogonal labeling techniques and cell-permeable dyes are required. In addition, the fluorophores should preferentially emit in the red spectral range to reduce potential phototoxic effects that can be induced by the STED light, which further restricts the choice of suitable markers. Here we demonstrate two-color STED imaging of living cells using various pairs of dyes that fulfill all of the above requirements. To this end, we combine click-chemistry-based protein labeling with other orthogonal and highly specific labeling methods, enabling long-term STED imaging of different target structures in living specimens.

## Introduction

Fluorescence microscopy has become an indispensable tool for cell-biological and biomedical research. Its ability to visualize cellular proteins of interest with high specificity and its applicability to living cells allow for the direct observation of cellular processes in their native environment. Due to the diffraction of light, the resolution of conventional fluorescence microscopes is limited [1] to around 250 nm, which is not sufficient for the detailed visualization of many tiny subcellular structures. The resolution limit has been overcome by super-resolution techniques, of which in particular stimulated emission depletion (STED) nanoscopy has been applied for live-cell imaging [2–4]. This technique uses the process of stimulated emission by a donut-shaped depletion beam to de-excite molecules in the outer regions of the excitation spot to obtain super-resolution images [5,6].

Many biological questions require simultaneous observation of more than one structure or feature. In conventional fluorescence microscopy, this can be achieved using dyes which are spectrally well separated. However, to ensure best performance and color co-alignment at nanometer scale, for STED microscopy it is advantageous to use two excitation lines together with a single STED laser [7,8]. For this, two fluorophores are required to be distinguished based on different excitation wavelengths while exhibiting similar emission wavelengths. In addition to that, fluorophores for STED need to be bright and photostable to obtain sufficient signal for multiple recordings at high resolution. Further, absorption and emission of the fluorophores should preferentially be in the orange to far-red spectral range. This has the advantage that light of lower energy can be used for excitation and depletion, which prevents phototoxic effects that could result from the excitation of cellular molecules by the STED light. In addition, light with a longer wavelength penetrates deeper into the sample and is less aberrated. All these requirements limit the choice of suitable labels for multi-color live-cell STED imaging.

Two different types of fluorophores are typically employed for live-cell labeling: fluorescent proteins (FPs) and organic dyes. Fluorescent proteins can be genetically fused to cellular proteins of interest and can be imaged after transfection without any additional labeling steps. Their main drawback for STED nanoscopy is their poor photostability, and also their comparatively low brightness in the red and far-red range [9]. On the contrary, organic fluorophores with high brightness and photostability are available throughout the full spectral range, but they require more complex labeling procedures than fluorescent proteins. To image the interior of living cells, the dyes need to be cell-permeable and must be bound specifically to the target protein, which can be achieved by different labeling techniques. For instance, the dye can be coupled with another organic molecule (hereafter referred to as the ligand) that binds to the target structure with high affinity (e.g., paclitaxel and its derivatives for the labeling of microtubules [10]). However, the use of such labels is restricted to the specific target structure, and for many cellular targets there are no suitable organic probes. Another more universal approach is a two-step labeling procedure in which a derivatized dye specifically reacts with a protein tag (e.g., SNAP-tag [11,12], CLIP-tag [13] or HaloTag [14]), which is in turn genetically fused to the protein of interest. One disadvantage of these self-labeling tags and fluorescent proteins is their large size, which can strongly influence the properties and behavior of the target protein. In addition, the resulting displacement of the fluorophore from the structure of interest impedes the optical separation of closely neighbored objects, which is of greatest significance if imaging is performed on the nanometer scale [15].

Another labeling strategy, which combines the advantages of the two before mentioned approaches, is the utilization of click chemistry. This method is based on the conjunction of two molecules by a highly selective chemical reaction and can be applied both *in vitro* and in living systems. For instance, it can be used for the detection of cellular synthesis of DNA upon its labeling with 5-ethenyl-2’-deoxyuridine (EdU) [16]. Protein labeling by click chemistry in living cells can be achieved by the incorporation of a modified amino acid during protein synthesis, which reacts in a second step with an organic fluorophore that is functionalized accordingly. Incorporation of the amino acid is most frequently realized by using the pyrrolysine system. This amino acid is used for protein synthesis only by certain archaea and bacteria where it is encoded by the amber codon UAG. Since this codon is relatively rare and does not encode any amino acid in most organisms, its usual function as a stop codon can be suppressed by the expression of pyrrolysyl-tRNA and pyrrolysyl-tRNA synthetase and the addition of pyrrolysine to the cells. By chemical modification of the pyrrolysine side chain, the system can be used to incorporate unnatural amino acids (uAAs) into the target protein in which a UAG codon was introduced at a suitable position (Fig 1A, B). For live-cell click labeling, the side chain of the amino acid can be functionalized with a strained trans-cyclooctene, which specifically reacts with a tetrazine group that is coupled with the fluorescent dye in a strain-promoted inverse electron-demand Diels-Alder cycloaddition (SPIEDAC) reaction [17] (Fig 1C).

**Figure 1.**
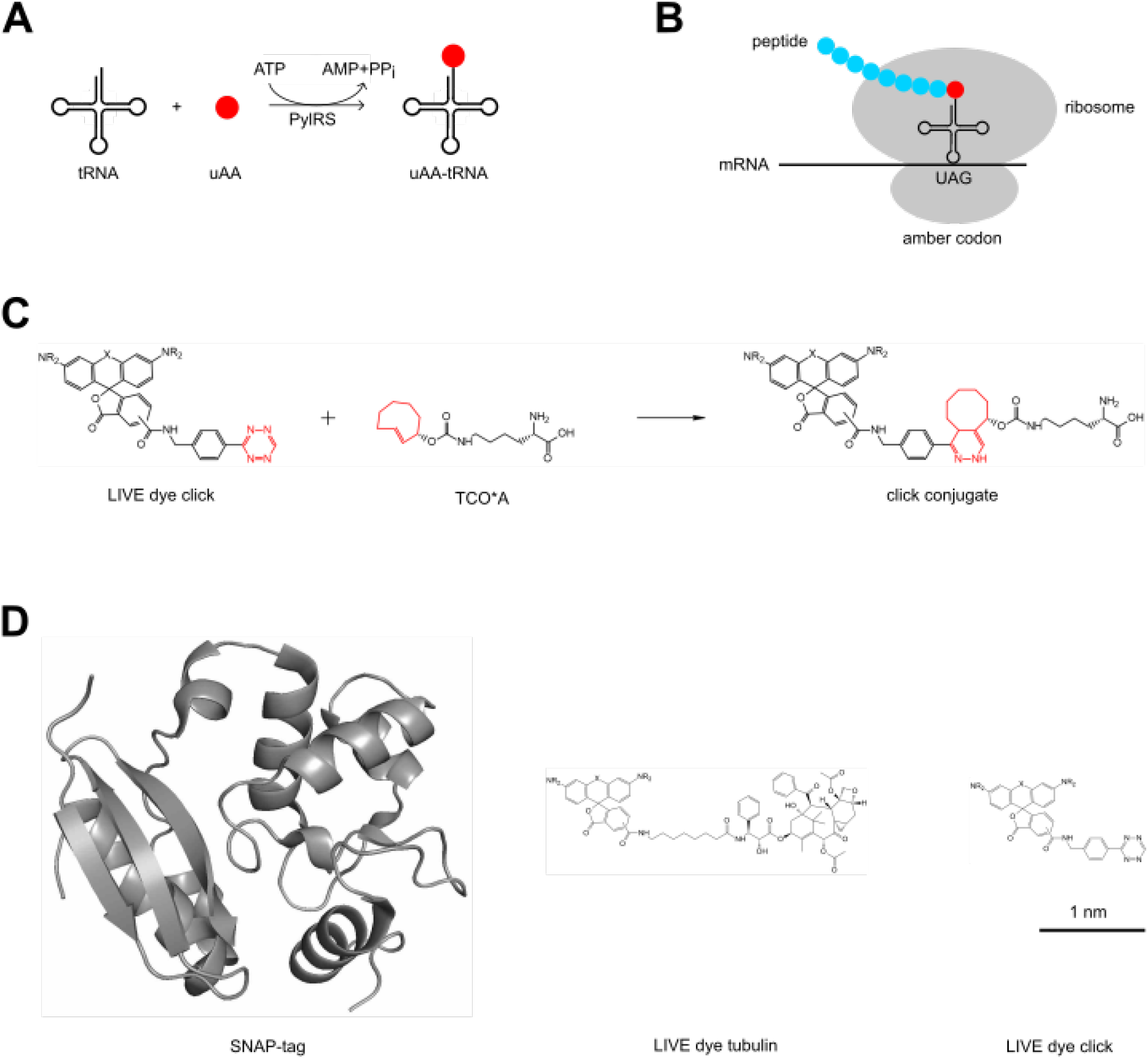
Click labeling of proteins in living cells. **(A)** PylRS catalyzes the activation of uAAs and their binding to a modified pyrrolysyl-tRNA. **(B)** The codon for these uAAs is the amber codon UAG, whose usual function as a stop codon is suppressed. **(C)** Click reaction between a tetrazine-functionalized rhodamine-based dye (LIVE dye click) and the strained trans-cyclooctene of the uAA TCO*A. **(D)** Size comparison of different labels for live-cell staining with organic fluorophores.

Click chemistry enables the site-specific labeling of proteins in living cells with tetrazine-functionalized fluorophores, which have a considerably smaller size than protein-based tags (Fig 1D). In addition, click labeling can be used together with other types of fluorescent labels and therefore facilitates the bioorthogonal labeling of different cellular targets with organic fluorophores. However, up to now only few dyes have been applied for intracellular click labeling of proteins in living cells [18], and of these only the silicon-rhodamine dye SiR [19] has been used for super-resolution imaging [18,20–22]. In this work, we have therefore tested tetrazine conjugates of several cell-permeable highly photostable organic fluorophores, which are suitable for confocal and STED imaging in the green to red spectral range, to increase the range of available dyes for live-cell click labeling. We further demonstrate two-color live-cell STED nanoscopy over extended time periods with low photobleaching.

## Results

### Photophysical properties of the dyes

The organic fluorophores we chose for tetrazine labeling are xanthene, carboxanthene or silicoxanthene dyes, which have been proven to have excellent cellular compatibility [23–27]. Their photophysical properties are shown in Table 1. LIVE 510 [23] is a green emitting fluorinated rhodamine dye that has been used for direct labeling via specific ligands to visualize microtubules, actin and mitochondria in living cells [24]. This dye has a high quantum yield and can be used for STED nanoscopy with a 595 nm depletion laser. The LIVE 460L fluorophore [25] is a long Stokes-shift dye that can be excited at 488 nm and emits in the range between 550 and 700 nm. Due to this strongly red-shifted emission compared to the absorption spectrum, this dye is well suited for STED nanoscopy with a 775 nm depletion laser and can be easily combined with other fluorophores for multi-color measurements. However, a potential problem of the LIVE 460L tetrazine probe is its low stability due to an intramolecular reaction. The rhodamine dye LIVE 550 and the two carbopyronins LIVE 590 and LIVE 610 are characterized by increased biocompatibility [26]. However, this feature is achieved only with one of the three possible chemical isomers. The different isomers of these orange and red emitting dyes have been tested on living cells and tissues by staining actin, microtubules, DNA, mitochondria and lysosomes as well as SNAP-tag and HaloTag proteins [26,28,29]. Due to a neighboring group effect in the isomer 4, cell permeability is increased with respect to the other two possible isomers. This results especially for direct labeling of the actin network, the microtubules and the DNA inside the nucleus in a high signal-to-background ratio when using sub-micromolar concentrations of isomer 4 probes instead of the other two possible isomer forms, respectively. Therefore, in case of the three dyes LIVE 550, LIVE 590 and LIVE 610, we used the chemical iosomer 4.

**Table 1.** Photophysical properties of the utilized dyes and fluorescent proteins. λ_abs. max_: absorption maximum; ε: extinction coefficient; λ_em. max_: emission maximum; τ: fluorescence lifetime; QY: quantum yield; ε∙QY: brightness (product of extinction coefficient and quantum yield). All data of the LIVE dyes were measured in PBS (pH = 7.4).

| Label | $\lambda_{\text{abs. max}}$<br>[nm] | $\epsilon$<br>[M<sup>-1</sup>·cm<sup>-1</sup>] | $\lambda_{\text{em. max}}$<br>[nm] | $\tau$<br>[ns] | QY | $\epsilon \cdot \text{QY}$<br>[M<sup>-1</sup>·cm<sup>-1</sup>] |
| --- | --- | --- | --- | --- | --- | --- |
| LIVE 510 [23] | 498 | 53,000 | 529 | 3.8 | 0.94 | 49,820 |
| LIVE 460L [26] | 455 | 28,000 | 617 | 2.0 | 0.17 | 4,760 |
| LIVE 550 [26] | 551 | 75,000 | 573 | 2.0 | 0.41 | 30,750 |
| LIVE 590 [26] | 585 | 105,000 | 609 | 3.5 | 0.60 | 63,000 |
| LIVE 610 [26] | 611 | 100,000 | 636 | 3.1 | 0.50 | 50,000 |
| mNeptune2 [40] | 599 | 89,000 | 651 | ND | 0.24 | 21,360 |
| mCardinal [40] | 604 | 87,000 | 659 | ND | 0.19 | 16,530 |
| mGarnet [36] | 598 | 95,000 | 670 | 0.8 | 0.09 | 8,550 |

### Live-cell STED nanoscopy of click-labeled actin

To apply the different tetrazine dyes for click labeling in living cells, we generated a plasmid encoding β-actin with amino acid 118 replaced by an amber codon (actin^K118TAG^) [20]. We chose the axial isomer of the uAA trans-cyclooct-2-ene-L-lysine (TCO*A) for integration into this construct due to its high acceptance by mutant tRNA synthetase, its high stability in mammalian cells and its fast reaction kinetic with tetrazines [30,31]. To incorporate the uAA into actin^K118TAG^, a modified tRNA and tRNA synthetase based on the pyrrolysine system were both expressed from a second plasmid. The double mutant Y271, Y349F of the pyrrolysyl-tRNA synthetase (PylRS) from *Methanosarcina barkeri* (MbPylRS^AF^) was used that efficiently reacts with non-natural lysine derivatives to form the corresponding aminoacyl-tRNAs [32,33]. To reduce the accumulation of MbPylRS^AF^ in the nucleus, we added a nuclear export signal (NES) at its N-terminus [34]. For improved incorporation of the uAA into the target protein, we additionally used the modified tRNA M15 which exhibits higher amber suppression efficiency due to an increased intracellular concentration [33].

For click labeling of actin, CV-1 cells were cotransfected with the plasmids containing actin^K118TAG^ and the tRNA/tRNA synthetase and incubated with TCO*A. After subsequent labeling with the tetrazine dye, cells were imaged using an abberior Instruments *STEDYCON* equipped with a top-stage incubation system to maintain the cells at 37 °C and 5% CO_2_. To compare the actin structure observed upon click labeling with an FP-based actin marker, we coexpressed EGFP-actin in cells labeled with actin^K118TAG^ and the tetrazine probe LIVE 610 click (S1 Fig). Both types of labels produced a highly similar actin staining, confirming the utility of the click-labeled actin^K118TAG^ construct for imaging of the native actin structure in living cells.

Next, we tested the performance of the dyes LIVE 610, 590, 550, 510 and 460L coupled to tetrazine amine for click labeling and live-cell STED nanoscopy (Fig 2, S2–S5 Fig). While labeling with LIVE 510 showed the anticipated actin pattern in confocal mode (S2 Fig), the probe was contrary to expectations not applicable for STED imaging with a 595 nm STED laser due to insufficient photostability. All other dyes were successfully applied for STED nanoscopy (Fig 2, S3–S5 Fig). Together, these results demonstrate that the cell-permeable probes LIVE 610, 590, 550 and 460L click can be used for click labeling of a TCO*A-containing protein of interest and STED imaging inside living cells.

**Figure 2.**
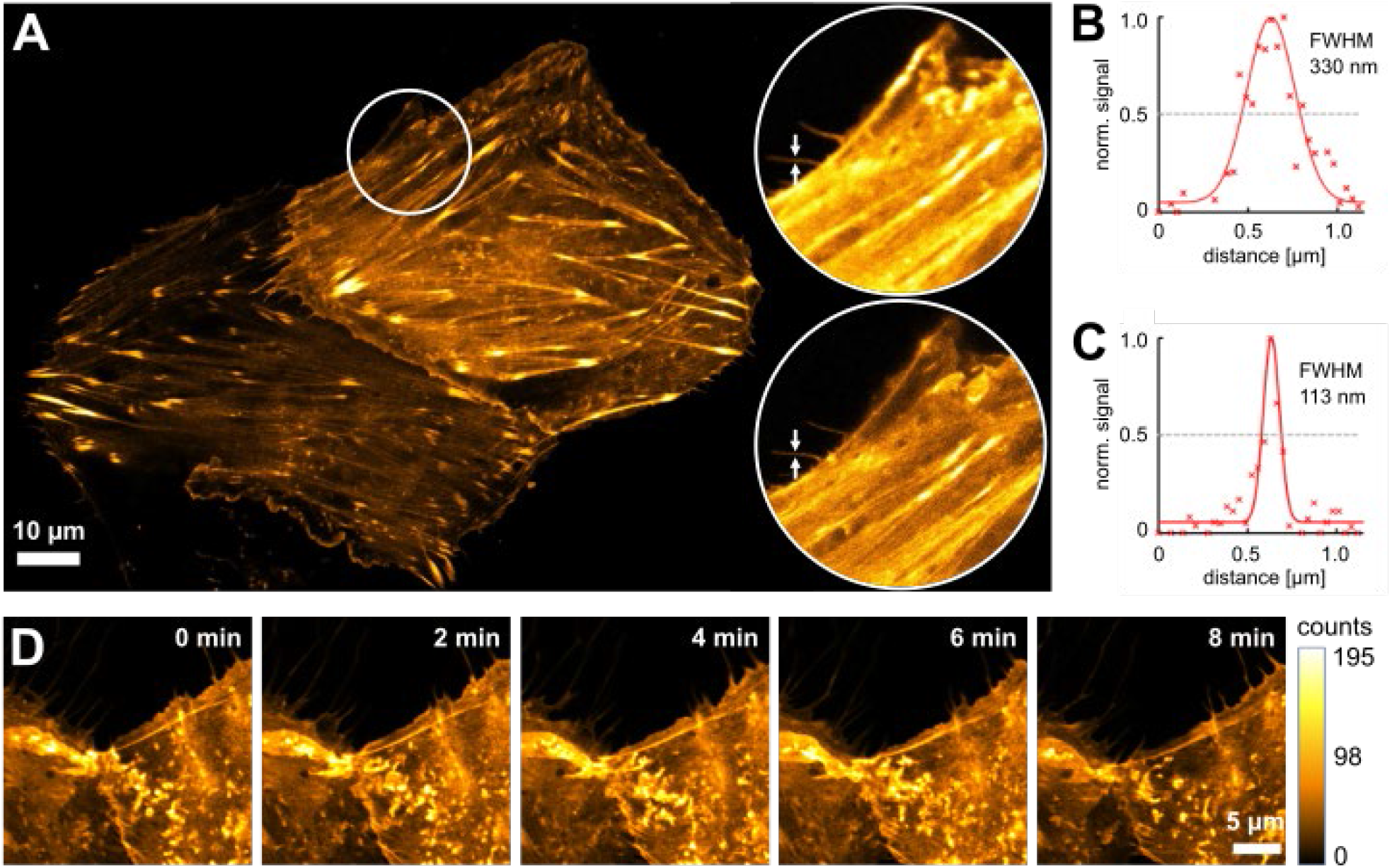
Fluorescence imaging of click-labeled actin filaments. **(A)** Living CV-1 cells expressing actin^K118TAG^ were labeled with TCO*A and LIVE 590 click. **(B, C)** The line profiles correspond to the filament marked with an arrow in the confocal (B) and STED (C) close-up images of (A), respectively. **(D)** Time series of actin filaments in living CV-1 cells stained with LIVE 590 click and measured in STED mode. During the time lapse, ten frames with an interval of 1 min were recorded. The fluorophore was excited with a 561 nm laser. For STED imaging, a 775 nm depletion laser was used.

### Time-lapse STED imaging with tetrazine dyes and fluorescent proteins

Specific labeling and imaging of proteins of interest in living cells are routinely performed using fluorescent proteins. In the far-red spectral range, the fluorescent proteins mNeptune2, mCardinal and mGarnet (Table 1) have been applied for STED nanoscopy [35,36] due to their relatively high brightness and photostability in comparison with other proteins in this wavelength range. However, due to the lower extinction coefficients and especially the much smaller quantum yields, their brightness (extinction coefficient multiplied by the quantum yield) is relatively low compared to the organic dyes in the same emission range. As a consequence, a higher excitation power might be needed for the fluorescent proteins, leading to higher cellular stress.

In order to compare the applicability of mNeptune2, mCardinal and mGarnet for long-term super-resolution microscopy with the LIVE dyes, we generated actin fusions of these proteins and performed time-lapse STED imaging in living cells. We chose LIVE 590 for comparison as this dye can be excited with the same 561 nm laser as the fluorescent proteins and used identical laser powers for all measurements. For LIVE 590, we performed click-labeling with actin^K118TAG^ and LIVE 590 click as described above. Whereas the fluorescence of LIVE 590 was only bleached to 50% of its initial value after ∼100 cycles of STED imaging, both mCardinal and mNeptune2 lost more than 50% of their fluorescence already after the first imaging cycle. mGarnet was more photostable than mCardinal and mNeptune2 and bleached to 50% of its maximum signal after ∼20 imaging frames (S6 Fig). However, the higher photostability of mGarnet is partly compromised by the low brightness of this protein, which is around 2-fold lower than that of mCardinal and mNeptune2 and only 14% of that of LIVE 590 (Table 1). Therefore, LIVE 590 is far superior to all fluorescent proteins tested both in terms of photostability and brightness, making it a more suitable label for STED nanoscopy of dynamic processes in living cells.

### Two-color long-term STED nanoscopy in living cells

Next, we applied different combinations of labels and dyes to perform two-color STED imaging of click-labeled actin filaments along with other cellular structures. For optimal spectral separation, the two dyes were excited at different wavelengths (561 or 640 nm, respectively) and their fluorescence signals were detected in different spectral windows that were chosen according to the respective fluorophore. For all measurements, the same 775 nm STED laser was used in order to keep photobleaching of the fluorophores and phototoxic effects at a minimum.

First, we combined click labeling with a SNAP-tag which can be used to label any cellular protein of interest with an organic dye (Fig 3A). For this purpose, we expressed the mitochondrial outer membrane protein OMP25 fused to a SNAP-tag and labeled it with LIVE 610 SNAP. Actin was labeled concomitantly with LIVE 550 click. STED imaging showed a significant improvement in resolution for both dyes when compared to confocal mode, allowing clear distinction of the mitochondrial membranes. Next, we used LIVE 550 click together with the cell-permeable probe LIVE 610 tubulin for simultaneous imaging of the actin and tubulin cytoskeleton in living cells (Fig 3B). The LIVE 610 tubulin probe consists of the carbopyronine dye LIVE 610 and the ligand cabazitaxel. In addition to high fluorogenicity, this probe shows good biocompatibility and high affinity for its target so that concentrations in the lower nM range are sufficient for complete labeling of the microtubules. Unlike click and SNAP-tag labeling, LIVE 610 tubulin therefore enables the direct labeling of tubulin without any genetic modification of the target protein. Likewise, we used the cholesterol-based probe STAR RED membrane to directly label the cell membrane for combined imaging of actin with LIVE 590 click (Fig 3C). Using cholesterol as membrane anchor and a linker between the dye and the anchor, the outside of the plasma membrane is specifically labeled and at the same time the dye is prevented from interacting with the membrane [37]. Both probe combinations showed specific labeling of the target structures and an increased resolution upon STED imaging. Hence, different types of labels may be combined with click labeling to perform two-color super-resolution microscopy in living cells with different pairs of organic dyes.

**Figure 3.**
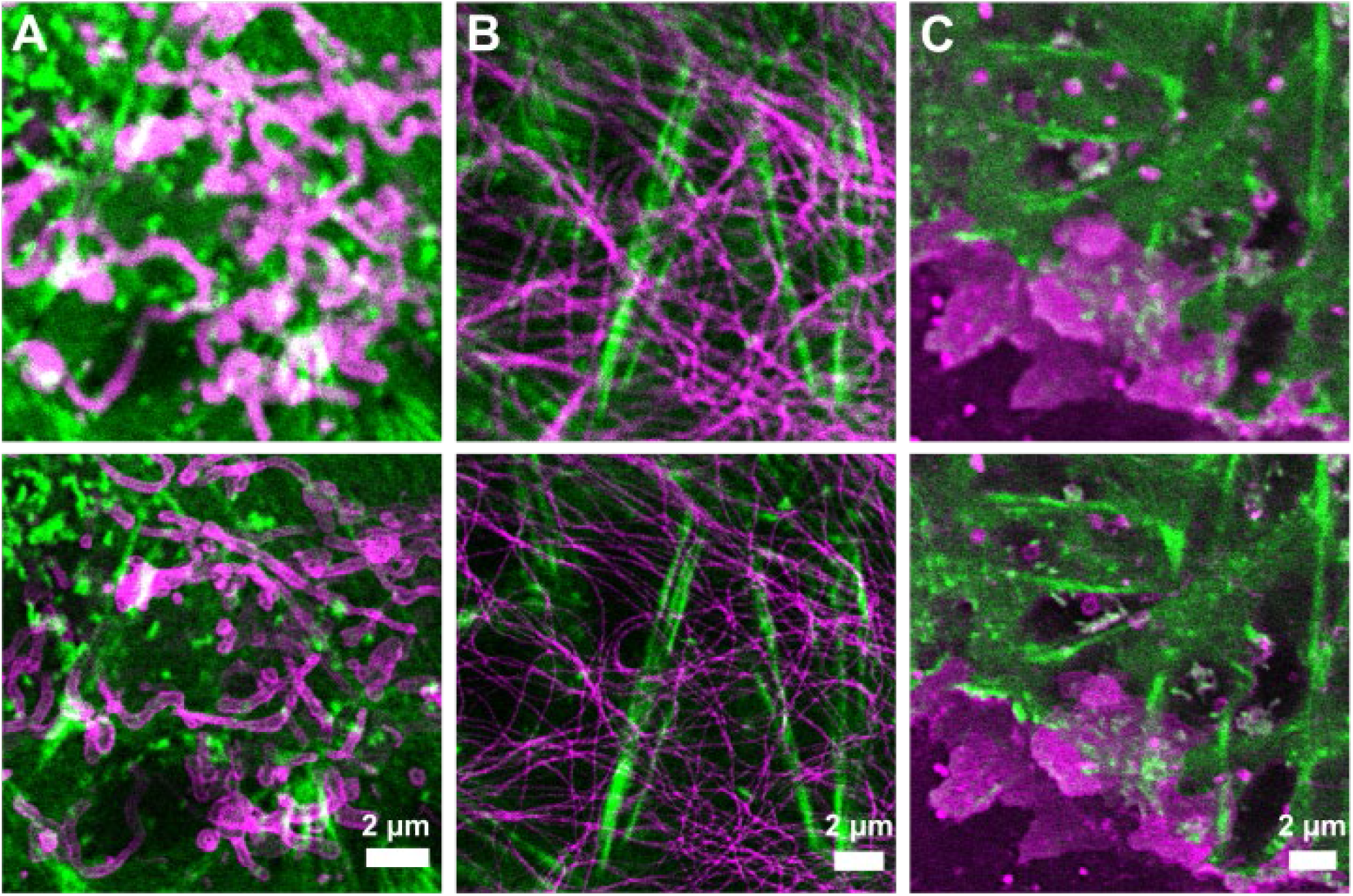
Two-color images of living CV-1 cells expressing actin^K118TAG^ incorporating TCO*A. Labeling was performed using the following combinations of fluorescent probes: **(A)** actin: LIVE 550 click, OMP25-SNAP: LIVE 610 SNAP, **(B)** actin: LIVE 550 click, microtubules: LIVE 610 tubulin, **(C)** actin: LIVE 590 click, plasma membrane: STAR RED membrane. The upper row represents confocal images while the bottom row shows the corresponding STED images. The fluorescent probes were excited with either a 561 nm or 640 nm laser. For the STED images, a 775 nm depletion laser was used.

For long-term STED imaging, we chose actin and OMP25 as intracellular targets and labeled them with LIVE 550 click and LIVE 610 SNAP, respectively (S1 Movie). Images were recorded every 30 s for a period of 30 min. The intact structure of the mitochondria and their movement in the cell throughout the entire observation time indicate that no major phototoxic effects were induced, confirming the good cellular tolerability of the far-red STED light. In addition, both dyes exhibit only moderate bleaching over the 60 recorded frames, demonstrating their usefulness for long-term STED measurements inside living cells.

## Discussion

STED nanoscopy has been applied for super-resolution live-cell imaging in different types of samples ranging from single-layered cell cultures to living animals [2,4]. To exploit the full potential of this technique, the development of suitable fluorophores and labeling procedures is of fundamental importance. While organic dyes are often more photostable than fluorescent proteins and therefore preferable for time-lapse STED measurements, the number of cell-permeable STED-compatible probes is still limited. We set out to expand the choice of available markers by combining several cell-permeable fluorescent dyes with SPIEDAC click labeling in living cells. By direct binding of the fluorophore to the protein of interest, this strategy enables the specific labeling of the target with the smallest possible label size, so that the influence of the label itself on protein function is kept at a minimum. In addition, with the recent development of improved super-resolution techniques such as MINFLUX [38] and MINSTED [15] that allow imaging at nanometer-scale resolution, the availability of small labels with minimal offset to the target is becoming increasingly important. This was demonstrated very recently by applying click labeling in MINFLUX nanoscopy whereby a localization precision of ∼2 nm was achieved in fixed cells [39].

By combining click labeling with several other live-cell-compatible probes, we demonstrated two-color STED nanoscopy of different target structures in living cells. The employed organic dyes emit in the orange/red region and exhibit higher photostability and brightness than fluorescent proteins in this spectral range. Hence, we successfully applied live-cell compatible dyes for long-term STED imaging using a single depletion laser at 775 nm which minimizes potential cell-damaging effects. By providing a new set of cell-permeable tetrazine probes, we expect that click labeling will become more widely used for super-resolution imaging inside living cells in the future.

## Methods

### Plasmid construction

A plasmid containing humanized PylRS from *M. barkeri* (MbPylRS) with the mutations Y271A and Y349F and the improved pyrrolysyl-tRNA M15 was generated based on the plasmid pNEU-hMbPylRS-4xU6M15 [33] (Addgene plasmid #105830). To reduce nuclear labeling [34], an NES was added to the PylRS by PCR with the primers NES-hMbPylRS NheI fwd and hMbPylRS NotI rev (for primer sequences, see S1 Table). The resulting PCR product was digested with NheI and NotI and ligated into pNEU-hMbPylRS-4xU6M15 digested with the same enzymes to produce pNEU-NES-hMbPylRS-4xU6M15.

To generate actin^K118TAG^ pcDNA3.1(+), human β-actin (ACTB) was PCR-amplified with the primers ACTB NheI fwd and ACTB BamHI rev. To insert the K118TAG mutation, a third primer ACTB K118TAG phosphorylated at its 5’ end and Taq DNA ligase (New England Biolabs) were added to the reaction mixture. The PCR product was cloned into pcDNA3.1(+) using the restriction enzymes NheI and BamHI.

For its N-terminal labeling with fluorescent proteins, β-actin was fused to EGFP, mNeptune2, mCardinal and mGarnet. For this purpose, ACTB was PCR-amplified using the primers ACTB BamHI fwd and ACTB NotI rev. The PCR product was cloned into pcDNA3.1(+) with BamHI and NotI. PCRs of the fluorescent proteins were performed with the following primers: EGFP: EGFP NheI fwd and EGFP HindIII rev, mNeptune2: mNeptune2 NheI fwd and EGFP HindIII rev, mCardinal: EGFP NheI fwd and EGFP HindIII rev, mGarnet: mGarnet NheI fwd and mGarnet HindIII rev. The PCR products were inserted into ACTB pcDNA3.1(+) with NheI and HindIII.

### Cell culture

CV-1 cells (Life Technologies) were cultured in DMEM (Gibco) with 4.5 g/l glucose supplemented with 10% FBS (Merck), 1 mM sodium pyruvate (Gibco), 100 units/ml penicillin and 100 μg/ml streptomycin (Merck). Cells were grown at 37 °C in 5% CO_2_.

### Live-cell labeling

For STED imaging, CV-1 cells were seeded on round coverslips (diameter 18 mm, thickness 0.16 mm, VWR) in 12-well plates in 1 ml of cell culture medium per coverslip. The following day, cells were transfected with a total amount of 1 µg DNA consisting of equal amounts of pNEU-NES-hMbPylRS_4xU6M15, actin^K118TAG^ pcDNA3.1(+) and optionally a construct containing either OMP25-SNAP-tag or EGFP-actin. For mNeptune2-, mCardinal- and mGarnet-actin, 1 µg DNA was used without any other plasmids. Transfections were performed using 2 µl jetPRIME transfection reagent (Polyplus-transfection) according to the manufacturer’s instructions. The medium was replaced by fresh cell culture medium 4 h post-transfection. TCO*A (Sirius Fine Chemicals SiChem GmbH) was added to the cells at a concentration of 200 µM.

The following day, cells were washed three times with cell culture medium to remove excess TCO*A and incubated at 37 °C and 5% CO_2_ for 1 h. Cells were then washed again and incubated with the tetrazine dye for click labeling of actin at a concentration of 2 µM for 30 min. Subsequently, the cells were washed and incubated in cell culture medium without phenol red for 30 min before imaging.

Labeling of OMP25-SNAP was performed with 1 µM LIVE 610 SNAP simultaneously with the application of the tetrazine dye. For labeling of the cell membrane, STAR RED membrane was applied in the final washing step at a concentration of 20 nM. Likewise, tubulin was labeled using LIVE 610 tubulin at a concentration of 0.2 µM together with 10 µM verapamil. Tetrazine dyes and other organic fluorescent probes were provided by Abberior GmbH.

### Live-cell STED nanoscopy

Confocal and STED microscopy images were acquired with a *STEDYCON* (Abberior Instruments GmbH). Pulsed lasers at 640 nm, 561 nm and 488 nm were used for fluorescence excitation. The detection windows were 650–700 nm for excitation at 640 nm and 575–625 nm for excitation at 561 nm. For excitation at 488 nm, detection windows of 575–625 nm and 505–545 nm were used for LIVE 460L and LIVE 510, respectively. For STED nanoscopy, a pulsed 775 nm STED laser (frequency 40 MHz, pulse length 1.1 ns) was additionally used. Single images were acquired with the following excitation powers: 0.20 mW at 488 nm, 0.16–0.17 mW at 561 nm and 0.13– mW at 640 nm. For STED imaging, the power of the 775 nm depletion laser was chosen according to the respective fluorophore (209 mW for LIVE 460L, 339 mW for LIVE 550, 194– 295 mW for LIVE 590, 82–101 mW for LIVE 610 and 45 mW for STAR RED). Time-lapse measurements of LIVE 590 and the red fluorescent proteins were performed with laser powers of mW at 561 nm and 222 mW at 775 nm. Laser powers for long-term two-color STED imaging (S1 Movie) were 0.16 mM at 561 nm and 157 mW at 775 nm for LIVE 550 and 0.13 mW and 63 mW at 775 nm for LIVE 610. The optical setup was connected to a wide-field IX83 fluorescence microscope (Olympus Europa SE & CO. KG) with a 60× oil immersion objective lens (NA = 1.40). Measurements were performed with a pixel dwell time of 10 µs, a pixel size of 30 nm and a line accumulation of 1.

After staining, cells were placed in an coverslip holder and covered with fresh cell culture medium without phenol red. For the time of measurement, cells were kept under ambient conditions (37 °C, 5% CO_2_, 90% relative humidity) by using a top-stage incubation system including the sample chamber H301-Mini (Oko-Lab).

The widths of actin filaments were identified by line profiles of the fluorescence intensities. The measurement points were interpolated using a Gaussian fit and its full width at half maximum (FWHM) was determined. The intensities of five adjacent pixels were averaged per measuring point.

## Acknowledgements

We thank Dr. Francesca Bottanelli (FU Berlin) for providing the OMP25-SNAP plasmid. pNEU-hMbPylRS-4xU6M15 was a gift from Irene Coin (Addgene plasmid #105830; http://n2t.net/addgene:105830; RRID:Addgene_105830).

## Supporting Information

**S1 Table.**
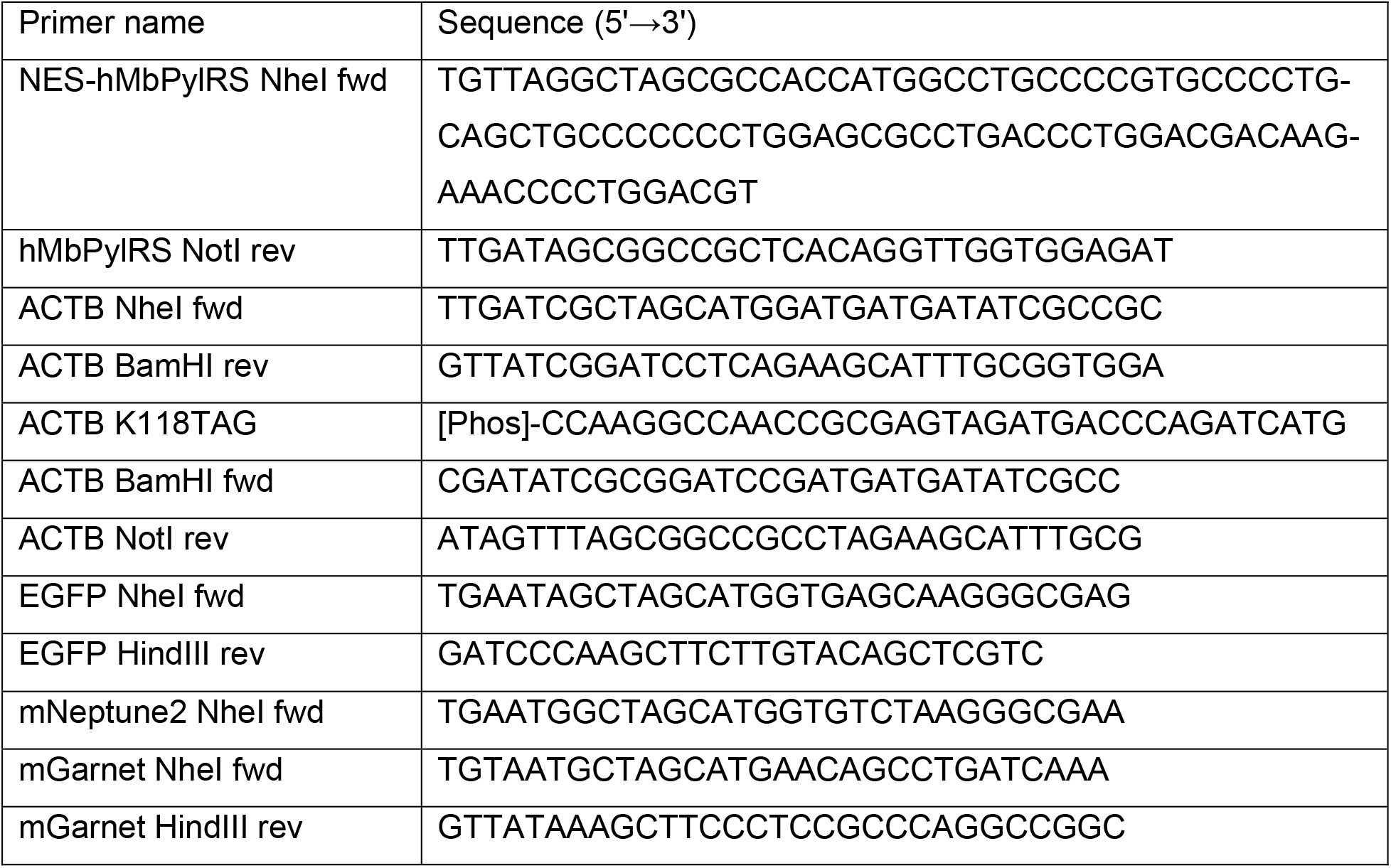
Primers used for plasmid construction.

**S1 Figure.**
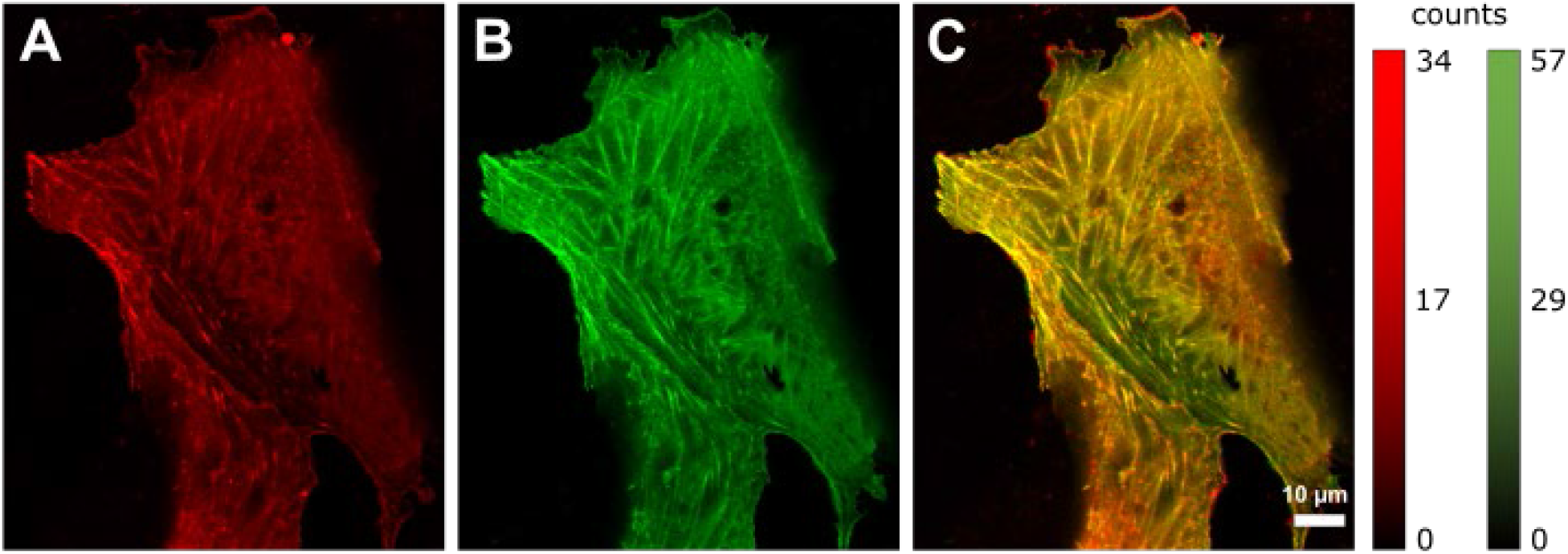
Actin structures of a living CV-1 cell. Cells were cotransfected with plasmids encoding EGFP-actin and actin^K118TAG^ and incubated with TCO*A. The cells were labeled with LIVE 610 click and imaged by confocal microscopy. **(A)** Fluorescence of LIVE 610 excited at 640 nm. **(B)** Fluorescence of EGFP excited at 488 nm. **(C)** Superposition of (A) and (B).

**S2 Figure.**
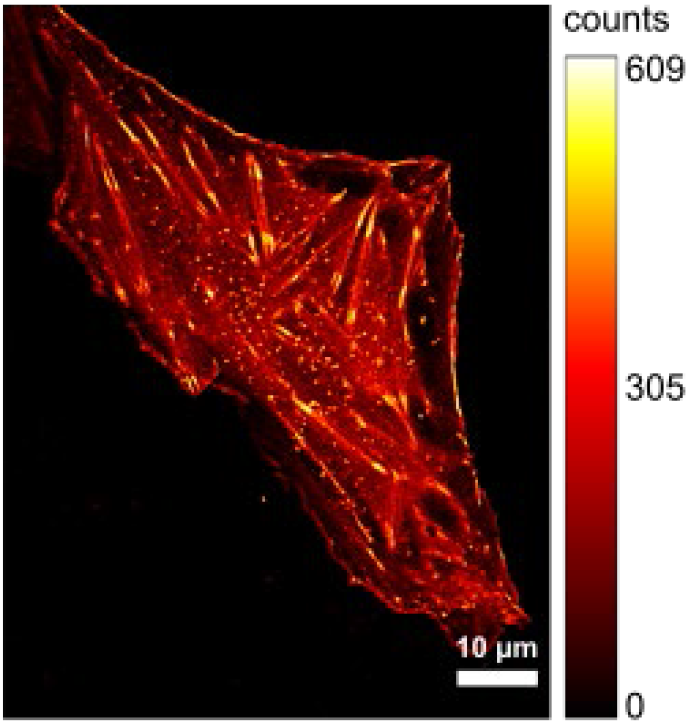
Fluorescence imaging of actin filaments click-labeled with LIVE 510. Confocal image of a living CV-1 cell expressing actin^K118TAG^ which was labeled with LIVE 510 click. The fluorophore was excited with a 488 nm laser.

**S3 Figure.**
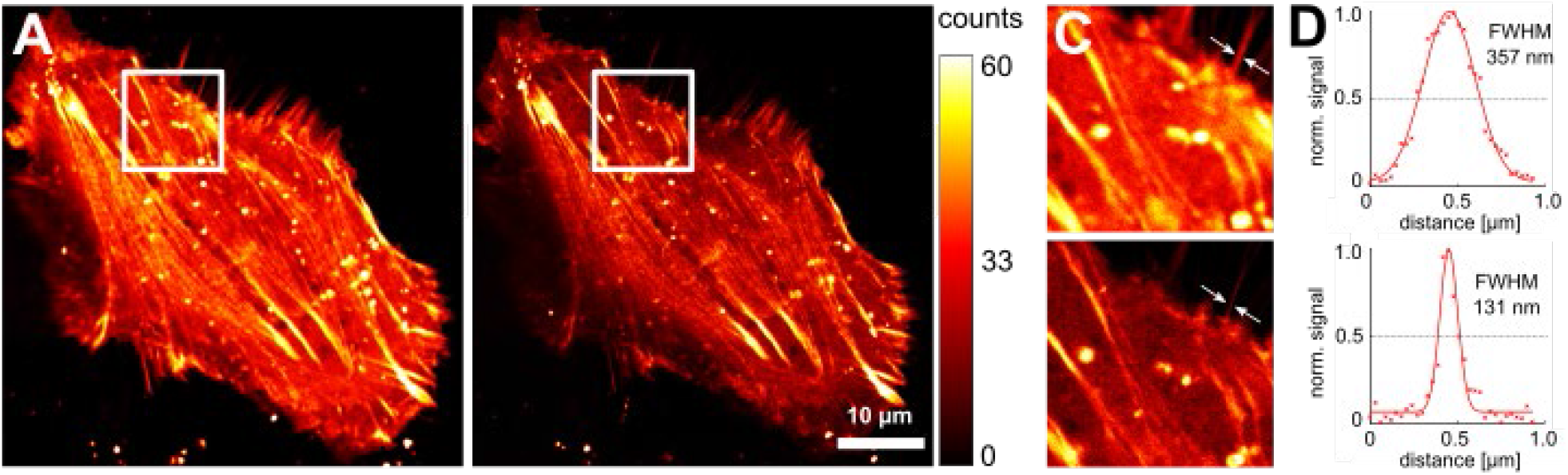
Fluorescence imaging of actin filaments click-labeled with LIVE 460L. **(A)** Confocal and **(B)** STED image of a living CV-1 cell expressing actin^K118TAG^ which was labeled with LIVE 460L click. **(C)** The upper panel shows a close-up of the confocal image (A), while the lower panel shows the close-up of the STED image (B). **(D)** Line profiles of the filament marked with an arrow in the confocal (upper panel) and STED image (lower panel) of (C).The fluorophore was excited with a 488 nm laser. For STED imaging, a 775 nm depletion laser was used.

**S4 Figure.**
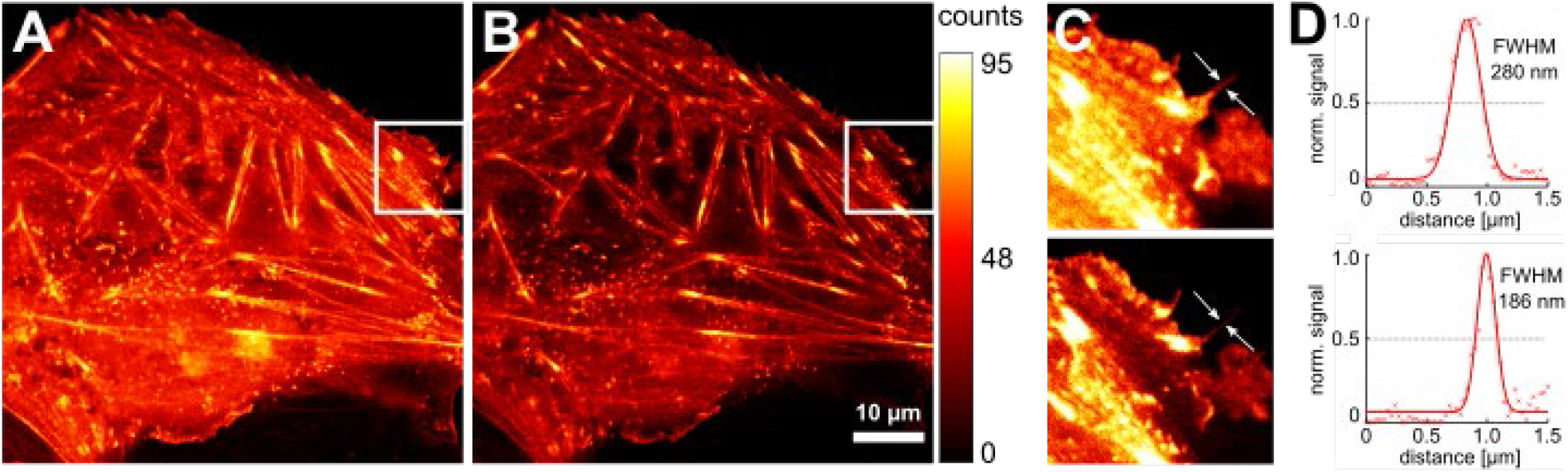
Fluorescence imaging of actin filaments click-labeled with LIVE 550. **(A)** Confocal and **(B)** STED image of a living CV-1 cell expressing actin^K118TAG^ which was labeled with LIVE 550 click. **(C)** The upper panel shows a close-up of the confocal image (A), while the lower panel shows the close-up of the STED image (B). **(D)** Line profiles of the filament marked with an arrow in the confocal (upper panel) and STED image (lower panel) of (C).The fluorophore was excited with a 561 nm laser. For STED imaging, a 775 nm depletion laser was used.

**S5 Figure.**
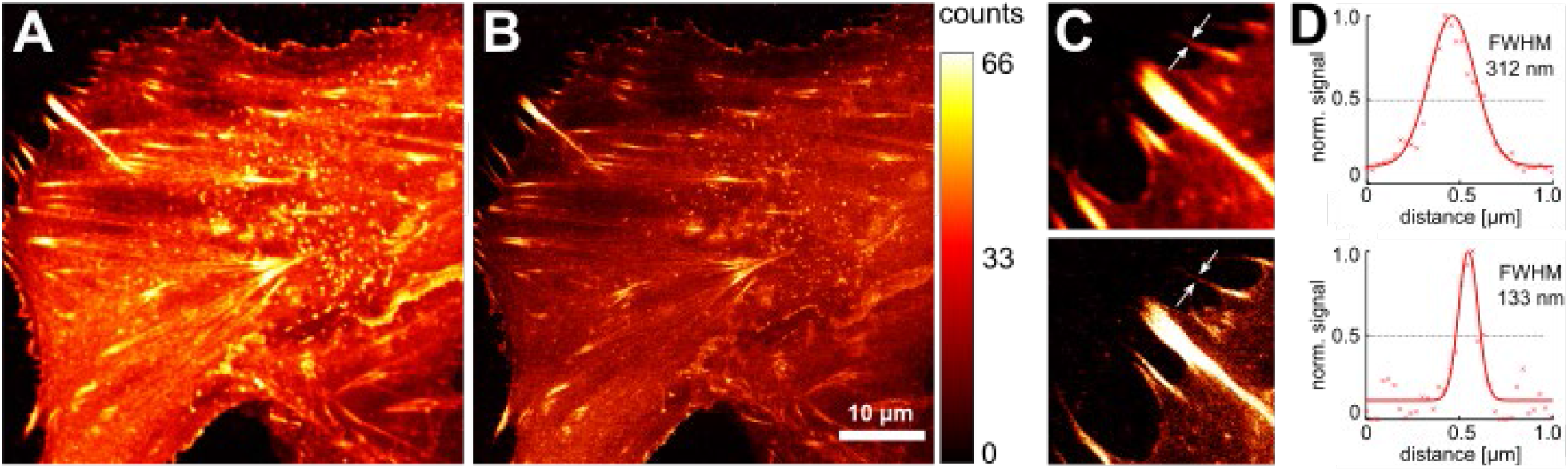
Fluorescence imaging of actin filaments click-labeled with LIVE 610. **(A)** Confocal and **(B)** STED image of a living CV-1 cell expressing actin^K118TAG^ which was labeled with LIVE 610 click. **(C)** The upper panel shows a close-up of the confocal image (A), while the lower panel shows the close-up of the STED image (B). **(D)** Line profiles of the filament marked with an arrow in the confocal (upper panel) and STED image (lower panel) of (C).The fluorophore was excited with a 640 nm laser. For STED imaging, a 775 nm depletion laser was used.

**S6 Figure.**
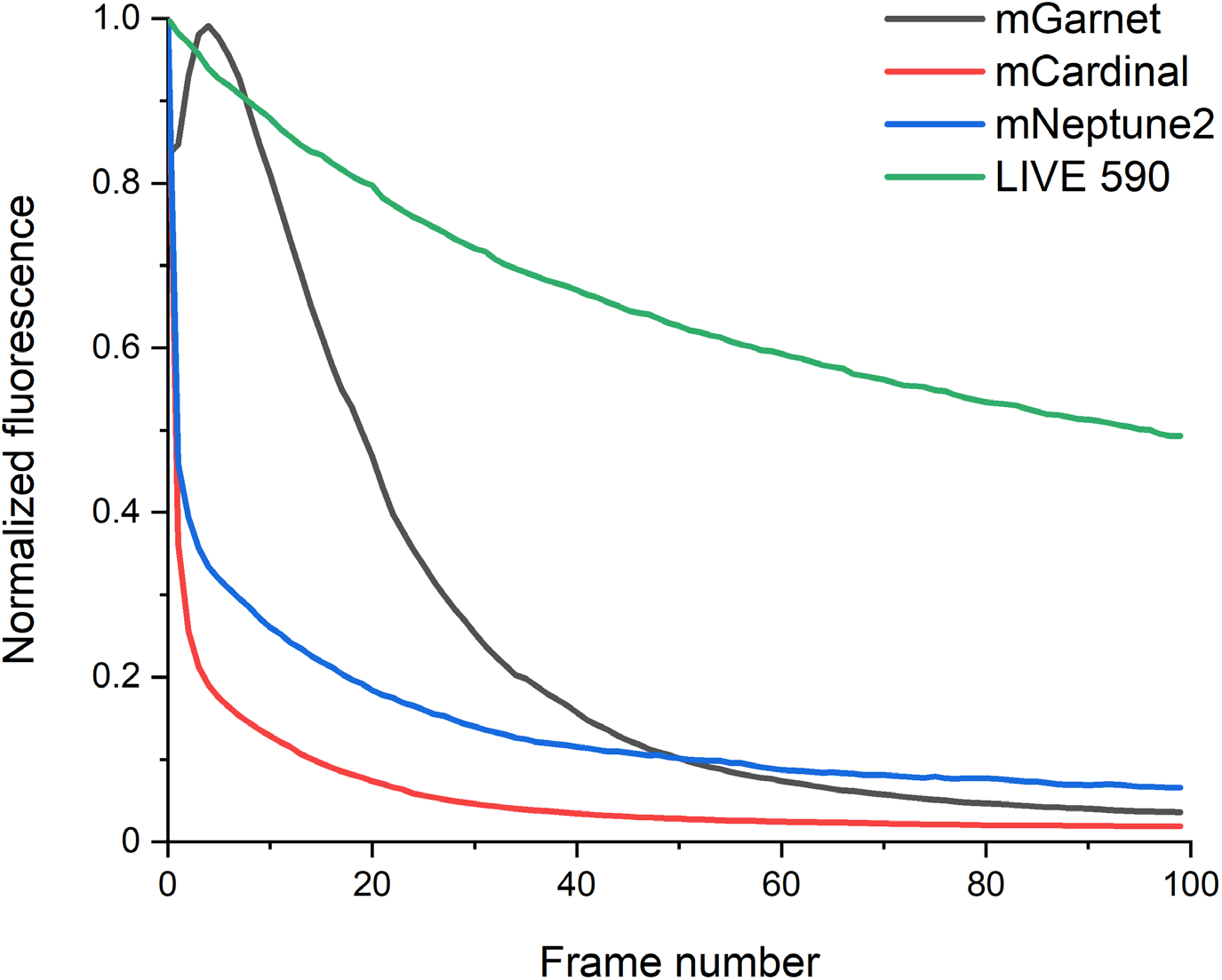
Bleaching of red fluorescent proteins and LIVE 590 during time-lapse STED imaging. The fluorescent proteins mGarnet, mCardinal and mNeptune2 were genetically fused to actin and expressed in CV-1 cells. For imaging of LIVE 590, CV-1 cells were transfected with actin^K118TAG^, incubated with TCO*A and labeled with LIVE 590 click. Imaging was performed in living cells using a 561 nm excitation laser and a 775 nm depletion laser with the same settings for all measurements. The values shown represent the average of three measurement series for each construct that were normalized to the maximum fluorescence signal.

**S1 Movie**. Two-color long-term STED imaging of actin and mitochondria. Living CV-1 cells expressing actin^K118TAG^ and OMP25-SNAP were incubated with TCO*A and labeled with LIVE 550 click and LIVE 610 SNAP. The dyes were excited with a 561 nm and a 640 nm laser, respectively, and depleted with a 775 nm STED laser. Images were recorded every 30 s for a period of 30 min.

## References

[1] Abbe E. Beiträge zur Theorie des Mikroskops und der mikroskopischen Wahrnehmung. Archiv f mikrosk Anatomie. 1873; 9:413–68. https://doi.org/10.1007/BF02956173.

[2] Hein B, Willig KI, Hell SW. Stimulated emission depletion (STED) nanoscopy of a fluorescent protein-labeled organelle inside a living cell. Proc Natl Acad Sci U S A. 2008; 105:14271–6. https://doi.org/-10.1073/pnas.0807705105.

[3] Pellett PA, Sun X, Gould TJ, Rothman JE, Xu MQ, Corrêa IR, et al. Two-color STED microscopy in living cells. Biomed Opt Express. 2011; 2:2364. https://doi.org/10.1364/BOE.2.002364.

[4] Berning S, Willig KI, Steffens H, Dibaj P, Hell SW. Nanoscopy in a living mouse brain. Science. 2012; 335:551. https://doi.org/10.1126/science.1215369.

[5] Hell SW, Wichmann J. Breaking the diffraction resolution limit by stimulated emission: stimulated-emission-depletion fluorescence microscopy. Opt Lett. 1994; 19:780–2. https://doi.org/10.1364/-OL.19.000780.

[6] Klar TA, Hell SW. Subdiffraction resolution in far-field fluorescence microscopy. Opt Lett. 1999; 24:954–6. http://dx.doi.org/10.1364/OL.24.000954.

[7] Schmidt R, Wurm CA, Jakobs S, Engelhardt J, Egner A, Hell SW. Spherical nanosized focal spot unravels the interior of cells. Nat Methods. 2008; 5:539–44. https://doi.org/10.1038/nmeth.1214.

[8] Göttfert F, Wurm CA, Mueller V, Berning S, Cordes VC, Honigmann A, et al. Coaligned dual-channel STED nanoscopy and molecular diffusion analysis at 20 nm resolution. Biophys J. 2013; 105:L01–3. https://doi.org/10.1016/j.bpj.2013.05.029.

[9] Stockhammer A, Bottanelli F. Appreciating the small things in life: STED microscopy in living cells. J Phys D: Appl Phys. 2021; 54:033001. https://doi.org/10.1088/1361-6463/abac81.

[10] Lukinavičius G, Reymond L, D’Este E, Masharina A, Göttfert F, Ta H, et al. Fluorogenic probes for live-cell imaging of the cytoskeleton. Nat Methods. 2014; 11:731–3. http://doi.org/10.1038/nmeth.2972.

[11] Juillerat A, Gronemeyer T, Keppler A, Gendreizig S, Pick H, Vogel H, et al. Directed evolution of O^6^-alkylguanine-DNA alkyltransferase for efficient labeling of fusion proteins with small molecules in vivo. Chem Biol. 2003; 10:313–7. https://doi.org/10.1016/S1074-5521(03)00068-1.

[12] Mollwitz B, Brunk E, Schmitt S, Pojer F, Bannwarth M, Schiltz M, et al. Directed evolution of the suicide protein O^6^-alkylguanine-DNA alkyltransferase for increased reactivity results in an alkylated protein with exceptional stability. Biochemistry. 2012; 51:986–94. https://doi.org/10.1021/bi2016537.

[13] Gautier A, Juillerat A, Heinis C, Corrêa IR, Kindermann M, Beaufils F, et al. An engineered protein tag for multiprotein labeling in living cells. Chem Biol. 2008; 15:128–36. https://doi.org/10.1016/-j.chembiol.2008.01.007.

[14] Los GV, Encell LP, McDougall MG, Hartzell DD, Karassina N, Zimprich C, et al. HaloTag: a novel protein labeling technology for cell imaging and protein analysis. ACS Chem Biol. 2008; 3:373–82. https://-doi.org/10.1021/cb800025k.

[15] Weber M, Leutenegger M, Stoldt S, Jakobs S, Mihaila TS, Butkevich AN, et al. MINSTED fluorescence localization and nanoscopy. Nat Photonics. 2021; 15:361–6. https://doi.org/10.1038/s41566-021-00774-2.

[16] Chehrehasa F, Meedeniya ACB, Dwyer P, Abrahamsen G, Mackay-Sim A. EdU, a new thymidine analogue for labelling proliferating cells in the nervous system. J Neurosci Methods. 2009; 177:122–30. https://doi.org/10.1016/j.jneumeth.2008.10.006.

[17] Liu F, Paton RS, Kim S, Liang Y, Houk KN. Diels-Alder reactivities of strained and unstrained cycloalkenes with normal and inverse-electron-demand dienes: activation barriers and distortion/interaction analysis. J Am Chem Soc. 2013; 135:15642–9. https://doi.org/10.1021/ja408437u.

[18] Arsić A, Hagemann C, Stajković N, Schubert T, Nikić-Spiegel I. Minimal genetically encoded tags for fluorescent protein labeling in living neurons. Nat Commun. 2022; 13:314. https://doi.org/10.1038/-s41467-022-27956-y.

[19] Lukinavičius G, Umezawa K, Olivier N, Honigmann A, Yang G, Plass T, et al. A near-infrared fluorophore for live-cell super-resolution microscopy of cellular proteins. Nat Chem. 2013; 5:132–9. https://-doi.org/10.1038/nchem.1546.

[20] Uttamapinant C, Howe JD, Lang K, Beránek V, Davis L, Mahesh M, et al. Genetic code expansion enables live-cell and super-resolution imaging of site-specifically labeled cellular proteins. J Am Chem Soc. 2015; 137:4602–5. 10.1021/ja512838z.

[21] Beliu G, Kurz AJ, Kuhlemann AC, Behringer-Pliess L, Meub M, Wolf N, et al. Bioorthogonal labeling with tetrazine-dyes for super-resolution microscopy. Commun Biol. 2019; 2:261. https://doi.org/10.1038/-s42003-019-0518-z.

[22] Werther P, Yserentant K, Braun F, Kaltwasser N, Popp C, Baalmann M, et al. Live-cell localization microscopy with a fluorogenic and self-blinking tetrazine probe. Angew Chem - Int Ed. 2020; 59:804–10. https://doi.org/10.1002/anie.201906806.

[23] Mitronova GY, Belov V, Bossi M, Wurm C, Meyer L, Medda R, et al. New fluorinated rhodamines for optical microscopy and nanoscopy. Chem - Eur J. 2010; 16:4477–88. https://doi.org/10.1002/-chem.200903272.

[24] Grimm F, Nizamov S, Belov VN. Green-emitting rhodamine dyes for vital labeling of cell organelles using STED super-resolution microscopy. ChemBioChem. 2019; 20:2248–54. https://doi.org/10.1002/-cbic.201900177.

[25] Butkevich AN, Lukinavičius G, D’Este E, Hell SW. Cell-permeant large Stokes shift dyes for transfection-free multicolor nanoscopy. J Am Chem Soc. 2017; 139:12378–81. https://doi.org/10.1021/-jacs.7b06412.

[26] Bucevičius J, Kostiuk G, Gerasimaitė R, Gilat T, Lukinavičius G. Enhancing the biocompatibility of rhodamine fluorescent probes by a neighbouring group effect. Chem Sci. 2020; 11:7313–23. https://-doi.org/10.1039/d0sc02154g.

[27] Soliman K, Grimm F, Wurm CA, Egner A. Predicting the membrane permeability of organic fluorescent probes by the deep neural network based lipophilicity descriptor DeepFl-LogP. Sci Rep. 2021; 11:6991. https://doi.org/10.1038/s41598-021-86460-3.

[28] Butkevich AN, Mitronova GY, Sidenstein SC, Klocke JL, Kamin D, Meineke DNH, et al. Fluorescent rhodamines and fluorogenic carbopyronines for super-resolution STED microscopy in living cells. Angew Chem Int Ed. 2016; 55:3290–4. https://doi.org/10.1002/anie.201511018.

[29] Bucevičius J, Gerasimaitė R, Kiszka KA, Pradhan S, Kostiuk G, Koenen T, et al. A general, highly efficient and facile synthesis of biocompatible rhodamine dyes and probes for live-cell multicolor nanoscopy. BioRxiv [Preprint]. 2022 bioRxiv 495683 [posted 2022 Jun 13; revised 2022 Jun 17; cited 2022 Jun 27]: [23 p.]. Available from: https://www.biorxiv.org/content/10.1101/2022.06.13.495683v2 doi: 10.1101/2022.06.13.495683

[30] Nikić I, Plass T, Schraidt O, Szymánski J, Briggs JAG, Schultz C, et al. Minimal tags for rapid dual-color live-cell labeling and super-resolution microscopy. Angew Chem - Int Ed. 2014; 53:2245–9. https://-doi.org/10.1002/anie.201309847.

[31] Hoffmann JE, Plass T, Nikić I, Aramburu IV, Koehler C, Gillandt H, et al. Highly stable trans-cyclooctene amino acids for live-cell labeling. Chem - Eur J. 2015; 21:12266–70. https://doi.org/10.1002/-chem.201501647.

[32] Yanagisawa T, Ishii R, Fukunaga R, Kobayashi T, Sakamoto K, Yokoyama S. Multistep engineering of pyrrolysyl-tRNA synthetase to genetically encode N^ε^-(o-azidobenzyloxycarbonyl) lysine for site-specific protein modification. Chem Biol. 2008; 15:1187–97. https://doi.org/10.1016/j.chembiol.2008.10.004.

[33] Serfling R, Lorenz C, Etzel M, Schicht G, Böttke T, Mörl M, et al. Designer tRNAs for efficient incorporation of non-canonical amino acids by the pyrrolysine system in mammalian cells. Nucleic Acids Res. 2018; 46:1–10. https://doi.org/10.1093/nar/gkx1156.

[34] Nikić I, Estrada Girona G, Kang JH, Paci G, Mikhaleva S, Koehler C, et al. Debugging eukaryotic genetic code expansion for site-specific click-PAINT super-resolution microscopy. Angew Chem - Int Ed. 2016; 55:16172–6. https://doi.org/10.1002/anie.201608284.

[35] Wegner W, Ilgen P, Gregor C, van Dort J, Mott AC, Steffens H, et al. In vivo mouse and live cell STED microscopy of neuronal actin plasticity using far-red emitting fluorescent proteins. Sci Rep. 2017; 7:11781. https://doi.org/10.1038/s41598-017-11827-4.

[36] Hense A, Prunsche B, Gao P, Ishitsuka Y, Nienhaus K, Nienhaus GU. Monomeric Garnet, a far-red fluorescent protein for live-cell STED imaging. Sci Rep. 2015; 5:18006. https://doi.org/10.1038/-srep18006.

[37] Mobarak E, Javanainen M, Kulig W, Honigmann A, Sezgin E, Aho N, et al. How to minimize dye-induced perturbations while studying biomembrane structure and dynamics: PEG linkers as a rational alternative. Biochim Biophys Acta Biomembr. 2018; 1860:2436–45. https://doi.org/10.1016/-j.bbamem.2018.07.003.

[38] Balzarotti F, Eilers Y, Gwosch KC, Gynnå AH, Westphal V, Stefani FD, et al. Nanometer resolution imaging and tracking of fluorescent molecules with minimal photon fluxes. Science. 2017; 355:606–12. https://doi.org/10.1126/science.aak9913.

[39] Mihaila TS, Bäte C, Ostersehlt LM, Pape JK, Keller-Findeisen J, Sahl SJ, et al. Enhanced incorporation of subnanometer tags into cellular proteins for fluorescence nanoscopy via optimized genetic code expansion. Proc Natl Acad Sci U S A. 2022; 119:e2201861119. https://doi.org/10.1073/pnas.2201861119.

[40] Chu J, Haynes RD, Corbel SY, Li P, González-González E, Burg JS, et al. Non-invasive intravital imaging of cellular differentiation with a bright red-excitable fluorescent protein. Nat Methods. 2014; 11:572–8. https://doi.org/10.1038/nmeth.2888.

